# Cumulative Cultural Evolution in Structured Populations

**DOI:** 10.64898/2026.04.15.718734

**Authors:** Rafael N. C. C. Leite, Sandro M. Reia, Alex Mesoudi, Paulo R. A. Campos

## Abstract

We extend previous models of cumulative cultural evolution by incorporating structured populations and social networks. We examine how connectivity and network topology shape the accumulation of cultural complexity under unbiased (copy randomly), indirectly biased (copy successful individuals), and directly biased (copy successful traits) transmission. We consider random, scalefree, and small-world networks, as well as the communication structures introduced by Mason and Watts, and derive analytical approximations for the homogeneous case. We find that the effects of social structure depend strongly on the form of transmission bias. Under unbiased transmission, network effects are weak except at very low connectivity. Under indirect bias, cultural complexity increases with connectivity, whereas direct bias shows optimal performance at intermediate connectivity, reflecting a trade-off between diffusion and diversity. Differences across topologies are generally modest once the average degree is fixed. Overall, our results show that no single social structure universally promotes cumulative cultural evolution; instead, its effects depend primarily on the dynamics of learning and innovation.

## I. INTRODUCTION

Cumulative cultural evolution (CCE) is the process by which innovations are socially transmitted, retained, recombined and progressively improved through social learning, making it central to human societies [1, 2]. Beneficial modifications are preserved and built upon over time, generating a ratchet effect that prevents the loss of successful innovations [3]. This allows populations to accumulate skills, technologies, and knowledge beyond what individuals can invent alone, helping explain the evolution of our species, ecological success, and adaptability [4]. Understanding the factors that promote or limit CCE remains a key research challenge. Recent work highlights demographic and social factors, such as population size, opportunities for social learning, and the reliability of cultural transmission [5, 6].

Early formal models, explicitly framed within an evolutionary perspective and inspired by population genetics and evolutionary biology, showed that larger effective populations can better maintain and accumulate complex cultural traits [7, 8]. Larger populations provide a larger pool of social learners, reduce the random loss of beneficial innovations, and increase the likelihood that rare improvements are retained over time. This outcome is corroborated by recent developments in agent-based and network-based modelling, which show that larger populations can host more innovators and reduce stochastic loss of useful traits [9–11], and is supported by archaeological [12], ethnographic [13] and experimental [14] evidence.

However, population size is only part of the picture [5]. CCE is also shaped by other demographic factors, such as the structure of social interactions. Connectivity and network architecture determine how innovations spread, persist, and recombine across a population. Although more connected populations may provide broader access to information and facilitate diffusion, experimental studies show that the effect of connectivity is not monotonic [15]. Fully connected populations may converge too quickly on the same solutions, thereby eroding the cultural diversity needed for exploration and recombination. On the other hand, fragmented or partially connected populations can harbor greater lineage diversity, which can lead to the development of complex solutions [16].

Cantor et al. [10] have shown that the relationship between social structure and CCE is highly contingent. In their simulations, the authors observed that no single network architecture consistently promotes CCE across all conditions. Indeed, architectures that favor the coexistence and recombination of cultural lineages in some circumstances may prevent or slow the subsequent diffusion of beneficial innovations in others. Their results further demonstrate that transmission dynamics are not a secondary ingredient but a central component of the process, and that distinct dynamics can lead to qualitatively different outcomes even on the same underlying social network. It is thus necessary to examine network topology and transmission dynamics jointly rather than in isolation. However, Cantor et al.’s [10] simulations used only one particular implementation of CCE (the ‘potions’ task [15]) in which traits are recombined across two semi-independent lineages. They also only implemented two simple forms of transmission bias (one-to-many and one-to-one), with random selection of demonstrators. There is a need to study both how social structure affects alternative models of CCE, and different forms of cultural transmission.

Our study builds on an alternative agent-based model of CCE introduced by Mesoudi [17], which explicitly represents cumulative culture as the sequential acquisition of functionally dependent cultural traits under limited effort budgets. This model incorporates three alternative forms of social transmission [18]: (i) unbiased transmission, where learners copy a randomly chosen demonstrator; (ii) indirectly biased transmission, where agents copy all of the traits of the individual with the highest overall payoff; and (iii) directly biased transmission, where individuals copy, at each functional level, the highest-payoff trait available in the previous generation. Mesoudi [17] showed that these three transmission rules generate markedly different outcomes for the degree of cultural complexity that can evolve, with direct bias typically supporting higher cultural complexity than indirect bias, and both outperforming unbiased transmission.

Here, we extend Mesoudi’s [17] framework to structured populations by embedding cultural transmission in social networks. Specifically, we investigate how three canonical network topologies–random, scale-free, and small-world networks–and different degrees of average connectivity affect CCE under each of the three transmission biases. We examine the conditions under which the effects of topology and connectivity manifest, and whether the impact of social structure is robust across transmission rules. Additionally, we also adopt the set of network topologies introduced by Mason [19, 20]. These networks provide a useful benchmark because they were designed to capture distinct patterns of connectivity and centralization while controlling for basic structural properties such as the number of nodes and edges. Using this family of networks allows us to isolate the effects of topology and assess how different communication structures which possess varying efficiency in spreading information shape the accumulation and transmission of cultural traits under the learning rules considered here.

## II. THE MODEL

In the original model [17], the population consists of *N* agents (*i* = 1, 2, *…, N*) that evolve over *T* discrete generations (*t* = 0, 1, *…, T* −1). A basic premise of the model is the existence of a functional hierarchy, in which each individual acquires a set of cultural traits that are functionally sequential: traits at level *s* must be acquired before traits at level *s* + 1.

Let 𝒳 = {1, *…, X*} denote the set of possible cultural traits. To each trait *x* ∈ 𝒳 one assigns a fixed fitness value *z*_*x*_. Only a small number of traits are highly effective, while the majority produce minimal or neutral impact. This is implemented by assigning fitness values drawn from an exponential distribution with parameter one, then squaring, multiplying by two, and rounding to the nearest integer, as proposed in Ref. [21]. Importantly, fitness depends only on the trait and not on the functional level at which it is acquired.

For a given individual *i*, let *x*_*i,s*_ ∈ 𝒳 denote the trait acquired at functional level *s*. The fitness of individual *i, Z*_*i*_, is then defined as the cumulative fitness of all acquired traits up to the highest functional level *s*_*max,i*_, i.e.,

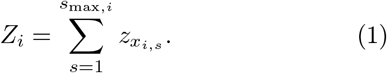

The mean population fitness is then used as a proxy for the cultural complexity of the population at generation *t*, and is calculated as

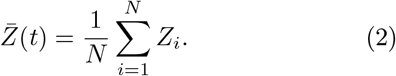

### A. Transmission rules

Every generation, the population is replaced by *N* individuals with no prior knowledge. Each individual has an effort budget *λ*, corresponding to the total effort they can allocate to learning cultural traits in their lifetime. The process of social learning begins by copying traits from the previous generation according to one of three transmission rules. Let 𝒫 (*t* − 1) = {1, *…, N*} denote the set of individuals in the previous generation.

In the first transmission model, dubbed indirect bias, a target *j*^*^ is selected by the new generation according to

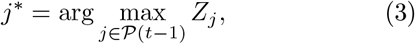

i.e., all individuals in the new generation copy the traits of the individual with the highest fitness in the previous generation. Copying each trait incurs a cost *c*_*s*_, measured in effort units, which are constant regardless of the trait’s fitness. The copying process continues until the learner either acquires all traits available to the target individual or exhausts their effort budget. If effort remains after copying, the learner engages in innovation. During this phase, a trait is randomly drawn from 𝒳 = {1, *…, X*} to occupy the next functional level *s*. If the trait is viable (i.e., its fitness is greater than zero), it is incorporated; otherwise, it is discarded, although the cost of innovation is still incurred. This process repeats until the effort budget is exhausted.

In the second transmission model, named direct bias, for each functional level *s*, the trait adopted by an individual of the new generation is

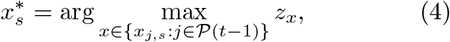

provided that at least one individual in the previous generation possesses a trait at level *s*. In other words, at each functional level *s*, individuals adopt the trait that yields the highest fitness among those available across every member of the previous generation. Once again, if effort still remains after the copying process, the innovation steps proceed, as previously described.

Both transmission rules can be considered plausible social learning strategies [18]. Imitating successful individuals, as in indirect bias, is a fast and low-cost way to acquire useful behavior, although it may also lead to the adoption of some less optimal traits. In contrast, direct bias is more likely to lead to the adoption of the most effective traits, but it generally requires more time and effort (although this greater cost is not explicitly modelled here).

In the third transmission model, unbiased transmission, a target *j*^*^ is drawn uniformly at random from 𝒫 (*t* − 1) with probability 1*/N*. Although this assumption is likely unrealistic as a description of how scientific and technological change occurs in practice, it serves as a useful benchmark for evaluating the effectiveness of the two biased transmission rules.

### B. Network topologies

While the original model assumes a homogeneous population in which every individual can potentially learn from every other, real social systems are typically structured by social ties that constrain who interacts with whom [22, 23]. We therefore extend the model to populations embedded in networks, in which nodes represent individuals and edges represent the set of possible social learning interactions.

To survey the influence of social architecture on CCE, we consider three well-studied network topologies: random, scale-free, and small-world networks. These topologies differ in key structural properties, particularly the degree distribution *P* (*k*), the clustering coefficient, and the average shortest path length, thereby allowing us to examine how patterns of connectivity shape cultural transmission and innovation.

In random networks, each pair of nodes is connected independently with probability *p*. The resulting degree distribution is given by a Poisson distribution in the limit of large *N*,

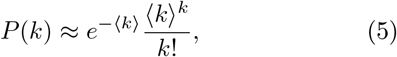

with average degree ⟨*k*⟩ ≈*p*(*N*− 1). This produces relatively homogeneous connectivity, low clustering, and the absence of highly connected hubs [24]. By contrast, scale-free networks (we use the Barabási-Albert version) are generated through growth and preferential attachment, leading to a broad degree distribution *P* (*k*) ∼ *k*^−*γ*^, with *γ* ≈ 3, strong degree heterogeneity, and the emergence of hubs that can disproportionately accelerate information spread [25, 26]. Finally, small-world networks interpolate between order and randomness. These networks preserve a narrow degree distribution, as in regular lattices, while combining high clustering with short average path lengths [27–29]. In our simulations, small-world networks were constructed by starting from rings and rewiring 10% of the connections. Additionally, we investigate the eight communication patterns proposed in Mason and Watts [19]. These networks all have the same size and density, namely *N* = 16 nodes and 24 edges, so that each node has degree three on average, but they differ markedly in their global organization, especially in average shortest path length 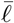 and in the distribution of betweenness centrality *B*_*i*_.

For each topology, we varied the average degree ⟨*k*⟩ to evaluate how local connectivity affects cultural evolution. In all simulations, population size took values *N* = 50, 100, and 300, the number of traits was fixed at *X* = 100, the individual effort budget at *λ* = 1000, and the cost parameters at *c*_*s*_ = 5 and *c*_*i*_ = 10. Each simulation tracked mean and maximum cultural complexity over time and results were averaged over 1000 independent runs.

## III. RESULTS

Before going through our simulation results, we first derive analytical predictions about the probability distribution of the traits’ payoffs, and then, in the following, we obtain some approximations for the cultural complexity of the population under unbiased and direct bias transmission models. These approximations consider homogeneous populations.

### A. Analytical derivations

#### 1. Derivation of the probability distribution of z_x_

Let *U* be a random variable whose probability density is *P*_*U*_ (*u*) = exp(−*u*). Then the probability density of variable *V* = *g*(*U*) = *U* ^2^ follows from

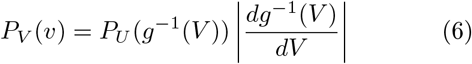

with 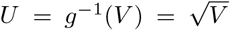. The resulting probability density *P*_*V*_ (*v*) equals 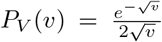. Similarly, it is straightforward to see that the variable *W* = 2*V* has probability density 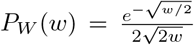. Its cumulative distribution function (CDF) is

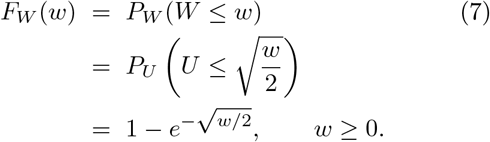

We now define a new discrete random variable *z*_*x*_ by rounding *W* to the nearest integer:

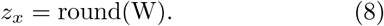

The support of *z*_*x*_ is

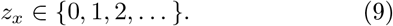

The event *z*_*x*_ = 0 occurs whenever

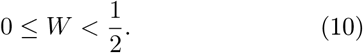

Therefore,

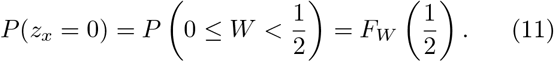

Using the expression for *F*_*W*_, we obtain

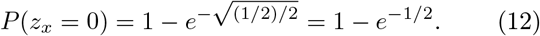

For any integer *n* ≥ 1, the event *z*_*x*_ = *n* occurs whenever

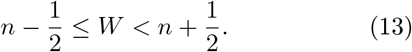

Hence,

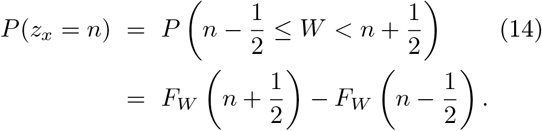

Substituting the cumulative distribution function of *W*, we find

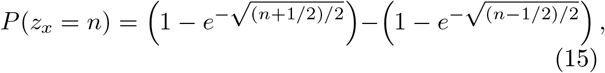

which simplifies to

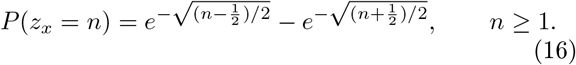

Thus, the probability distribution of *z*_*x*_ is

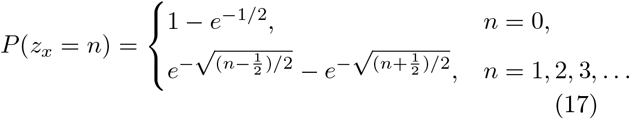

In conclusion, *z*_*x*_ is a discrete nonnegative integer-valued random variable obtained by rounding the continuous variable *W* = 2*X*^2^, and its distribution is completely determined by the expression above. It is useful to compute the expected value of *W*, since this allows us to estimate the expected complexity of the unbiased model. Because non-viable traits (null fitness) are not incorporated by individuals, we are mainly interested in estimating

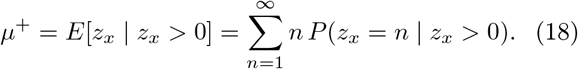

which equals

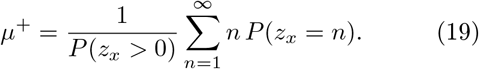

Using our derivations in Eq. (17), numerically we obtain that

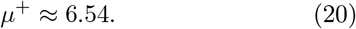

In the stationary regime, the highest function level is 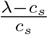, and so the cultural complexity for the unbiased model becomes

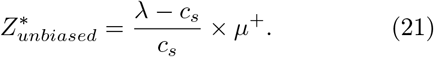

#### 2. The direct bias case

An approximation can be derived for the direct bias transmission model. In the approximation, one considers a non-structured population of size *N*, and for the derivation, one considers that at each level *s, N* independent draws of traits among *X* available traits are performed. The relevant number of candidate traits is not the total repertoire size *X*, but the expected number of distinct traits represented in a sample of *N* draws. This quantity is [30]

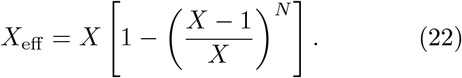

As a second approximation, the stationary maximum complexity under direct bias may then be estimated as

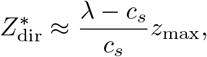

where *z*_max_ is the largest payoff effectively available in the trait repertoire of size *X*_eff_. In this case,

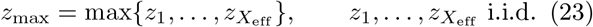

At this point, we will ignore the small correction due to rounding, such that

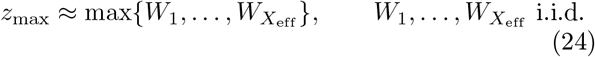

The typical maximum *W*_max_ among *X*_eff_ draws is obtained from

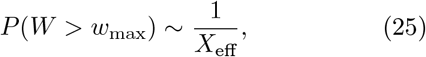

that is,

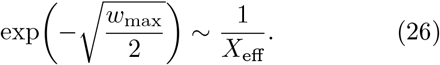

Therefore,

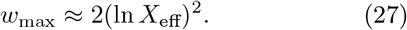

As we ignored corrections due to rounding, *z*_max_ equals *w*_max_, and so

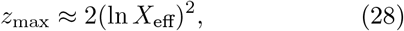

which yields the stationary estimate

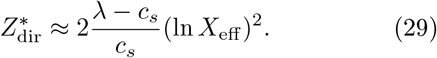

A more refined approximation, conditioning on viable traits, is found by replacing *X*_eff_ with *X*_eff_ */P* (*W >* 1*/*2). In this case,

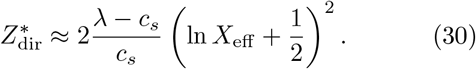

#### 3. The indirect bias case

Unfortunately, we could only obtain a poor estimate of the mean cultural complexity at stationarity. In the indirect bias transmission model, the complexity itself is selected at each round. But the complexity of each individual is just the sum of *s*_*max*_ variable *z*_*x*_. As we know, it means *µ*^+^ = 6.54, and its variance, 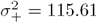, and following the Central Limit Theorem [31], the resulting distribution of *Z*_*j*_ is given by a Gaussian distribution of mean 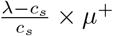 and variance 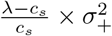. Following the same reasoning, as previously, we want to estimate *Z*_max_ such that

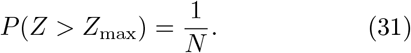

But *P* (*Z > Z*_max_) = 1 − *F*_*Z*_(*Z*_max_), where *F*_*Z*_(*Z*_max_) is the cumulative function distribution of *Z*. As we know the mean and variance of *Z*, so *Z*_max_ is the numerical solution of

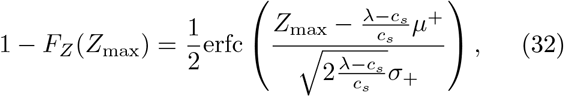

where erfc(*x*) is the complementary error function. The reason the prediction above underestimates the population complexity at stationarity is that *Z*_max_ itself undergoes optimization steps before stationarity is reached. Therefore, the solution of Eq. (32) works as a lower bound estimate of 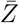 at stationarity.

### B. Simulation results

We begin by examining how network topology and average connectivity affect CCE under the three transmission rules. Figure 1 shows the temporal evolution of mean cultural complexity, 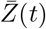, for representative values of the average degree ⟨*k*⟩ in random, scale-free, and small-world networks. As already noted in Ref. [17] and demonstrated in the previous section analytically, dynamics depend strongly on the transmission rule. Under unbiased transmission, the trajectories are nearly insensitive to ⟨*k*⟩ and to the network class, except for random networks at very low connectivity (⟨*k*⟩ = 2), where the mean cultural complexity settles at a lower level. At this low degree of connectivity, the graph is very likely not to comprise a single component. Thus, one has a set of unconnected components, thereby reducing the effective population size. Otherwise, network class and connectivity have no effect.

**Figure 1.**
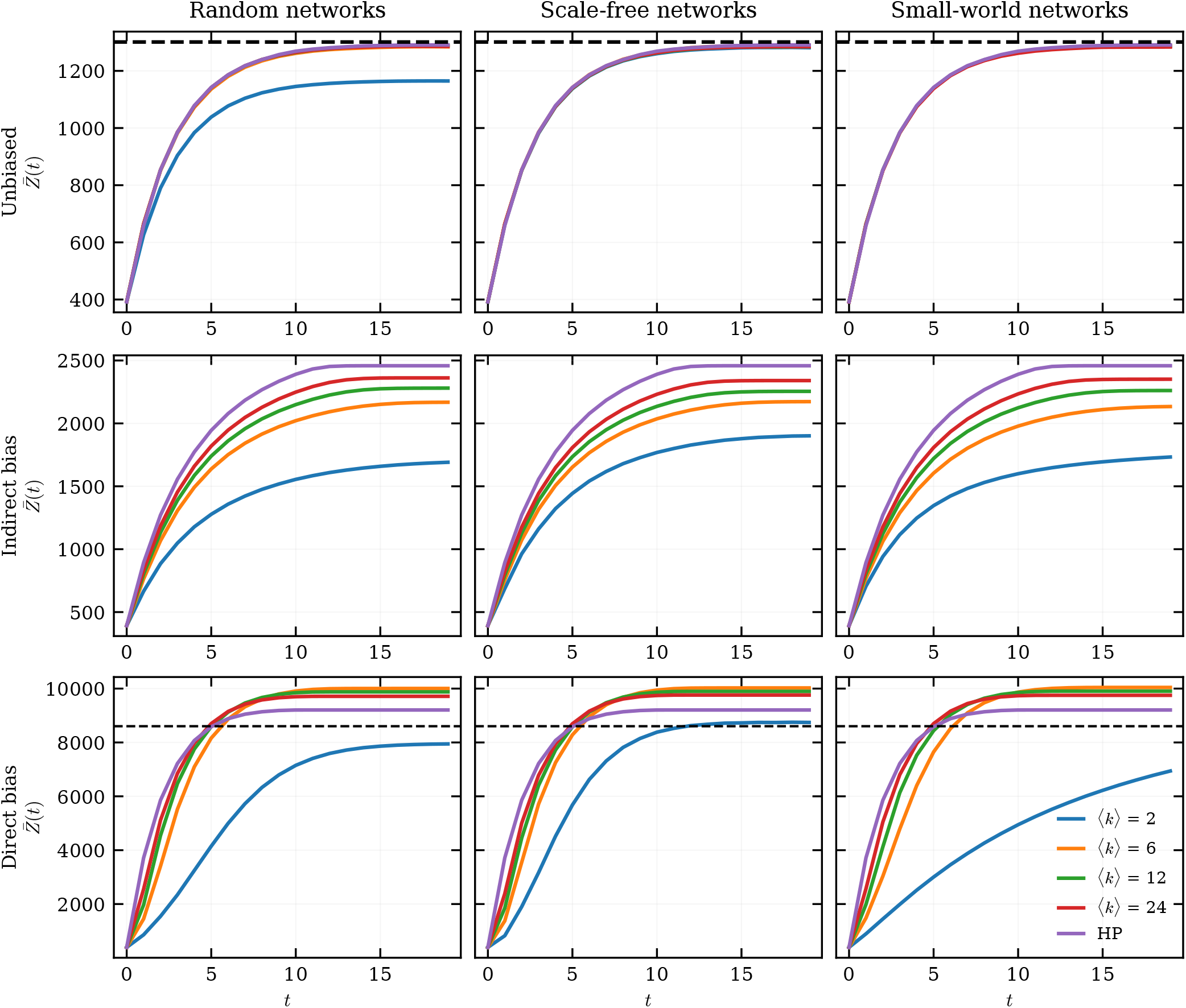
Temporal evolution of mean cultural complexity for representative values of average connectivity in random, scale-free, and small-world networks for each transmission rule. Parameter values are *N* = 100, *λ* = 1000, *c*_*s*_ = 5 and, *c*_*i*_ = 10. Horizontal dotted lines for unbiased and directly biased transmission show analytically derived expected equilibrium cultural complexity. HP = homogeneous populations.

By contrast, both biased transmission rules exhibit a clearer dependence on the mean connectivity. In the indirect bias case, increasing ⟨*k*⟩ results in systematically higher values of the mean complexity 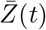 and a faster approach to equilibrium. In turn, the direct bias transmission model displays a different scenario, where the dependence of 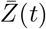 on mean connectivity is no longer monotonic, but instead peaks at intermediate ⟨*k*⟩. In both cases, differences among the three network topologies remain comparatively modest.

Figure 1 also shows results for homogeneous populations (HP), in which individuals can socially learn and interact with every other individual in the population. The dashed-lines present our analytical approximations for the stationary situation and the homogeneous case for the unbiased and directly biased transmission models. For unbiased transmission we obtain excellent agreement between simulations and the theoretical prediction (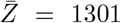 for these parameter values). For direct bias our analytical derivation 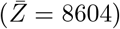 underestimates 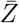. Though at larger *N*, for instance when *N* = 300 (see Figure S2), there is a perfect match between simulations and our analytical prediction. Therefore, for large *N*, the theoretical prediction, Eq. 30, is accurate. For indirect bias, as previously noted, the analytical approximation underestimates 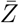.

For instance, the numerical solution of Eq. (32) predicts 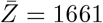 for *N* = 100 and 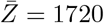 for *N* = 300, whereas for a homogeneous (unstructured) population, the simulation results in Figure 1 yield 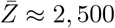.

Overall, the outcomes in Figure 1 indicate that connectedness is a primary determinant of temporal dynamics, whereas other structural properties of the networks do not appear to manifest. A relevant aspect of the findings in Figure 1 is that, for direct bias, the structured populations outperform the unstructured homogenous population, whether the latter is derived analytically or via simulation. Direct bias selects, at each functional level, the highest-fitness trait available in the local neighborhood, and thus leads to slower convergence and, consequently, homogenization of the population at higher fitness traits.

To assess how connectivity affects the final level of cultural accumulation, Figure 2 presents the maximum cultural complexity attained as a function of ⟨*k*⟩ for the three network families and for different population sizes. The unbiased rule exhibits comparatively weak dependence on connectivity, except in the low-connectivity regime, where random graphs exhibit a spurious effect arising from unconnected components. Under the indirect bias rule, the cultural complexity at stationarity rises monotonically with ⟨*k*⟩ across all three topologies. A different scenario arises under the direct bias model, in which the maximum cultural complexity is a one-humped function of ⟨*k*⟩. This feature is observed across the three topologies.

**Figure 2.**
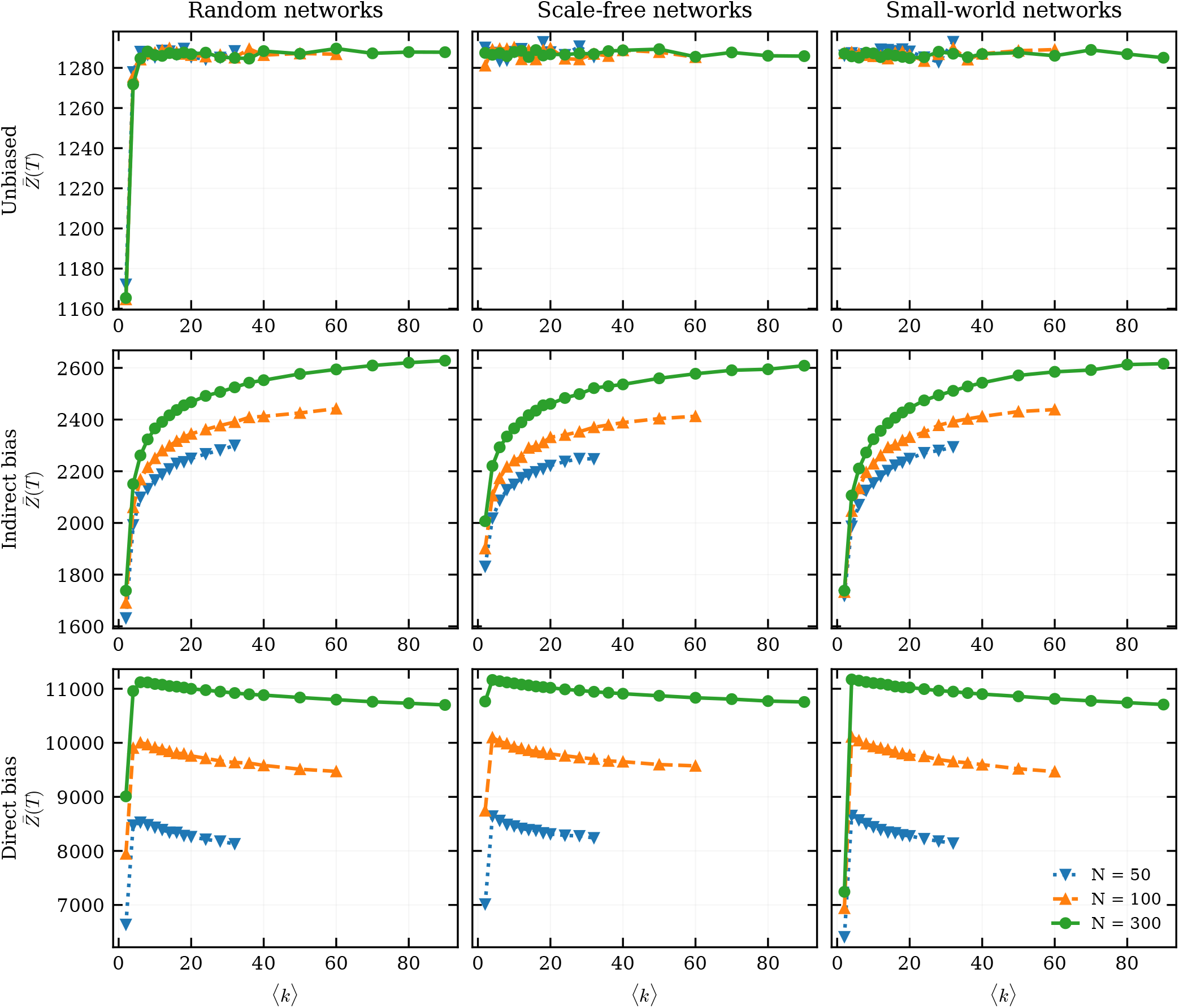
Maximum cultural complexity as a function of connectivity ⟨*k*⟩ and population size *N* in random, scale-free, and small-world networks for each transmission rule. Parameter values are *λ* = 1000, *c*_*s*_ = 5, and *c*_*i*_ = 10.

The existence of an optimal level of connectivity highlights that a trade-off occurs between at high connectivity the prompt diffusion of the currently best available traits and rapid population homogenization, albeit with a reduction in cultural diversity and reduced scope for further innovation, and at low connectivity the slower diffusion of traits, albeit with a higher level of diversity [16]. In the same plot, the role of demography is explored, and, as expected from previous studies [7–10], a larger population leads to higher mean population complexity across all scenarios, except for the unbiased model, in which the target individual for all individuals is a single randomly chosen agent from a previous generation. Because there is no underlying maximization principle, the outcome is unsurprising.

Finally, Figure 3 characterizes the speed of convergence by showing the equilibrium time as a function of ⟨*k*⟩. Here, the equilibrium time is defined as the earliest timestep at which the relative variation of 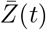 within a sliding window of four generations falls below 1%. Under unbiased transmission, the time to equilibrium is nearly constant, remaining close to 11 generations and showing little sensitivity to either connectivity or topology. The scenario is distinct under both direct and in-direct bias rules, where we see a prominent effect of ⟨*k*⟩ on equilibrium time. At low connectivities, both rules converge more slowly than the unbiased model, and as connectivity increases, their equilibrium times decrease monotonically. This effect is more pronounced for the direct bias rule, which converges fastest at high ⟨*k*⟩, attaining equilibrium in roughly 8 generations, ultimately faster than unbiased transmission. Indirect bias levels off at somewhat larger times, and remains slower than unbiased transmission even at high ⟨*k*⟩. As in the previous figures, differences among random, scale-free, and small-world networks are present but secondary relative to the effect of the transmission rule itself. This corroborates the observation above that direct bias is disadvantaged at high connectivity due to its speed, with sub-optimal traits being selected and diffusing prematurely.

**Figure 3.**
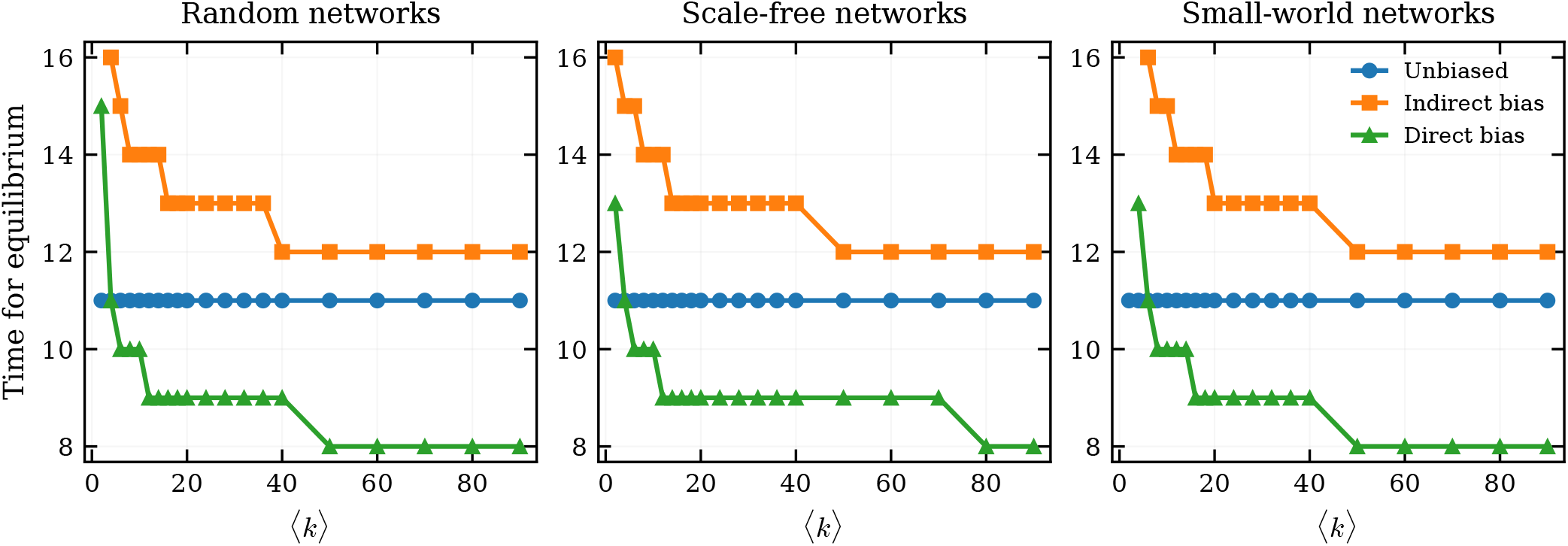
Time for equilibrium as a function of network connectivity ⟨*k*⟩in random, scale-free, and small-world networks for each transmission rule. Parameter values are *N* = 100, *λ* = 1000, *c*_*s*_ = 5, and *c*_*i*_ = 10.

### C. Communication structure in the Mason–Watts networks

To further examine how communication patterns influence CCE, we examine eight distinct network topologies illustrated in Figure 4. All networks contain 16 nodes and 24 edges, enabling regulated comparisons in which only structural organization varies [19]. Each of these eight networks is designed to present a clear structural purpose. Network A minimizes average betweenness centrality, thus mitigating bottlenecks on information flow. Network B minimizes average clustering. Network C maximizes maximum closeness centrality. Network D maximizes variance in structural constraints, resulting in heterogeneous local connectivity. The remaining four networks are less efficient at information propagation. Network E maximizes average clustering. Network F maximizes maximum betweenness, concentrating information flow in a few nodes. Network G minimizes maximum closeness, limiting extreme centralization. Network H maximizes average betweenness, increasing reliance on intermediary nodes.

**Figure 4.**
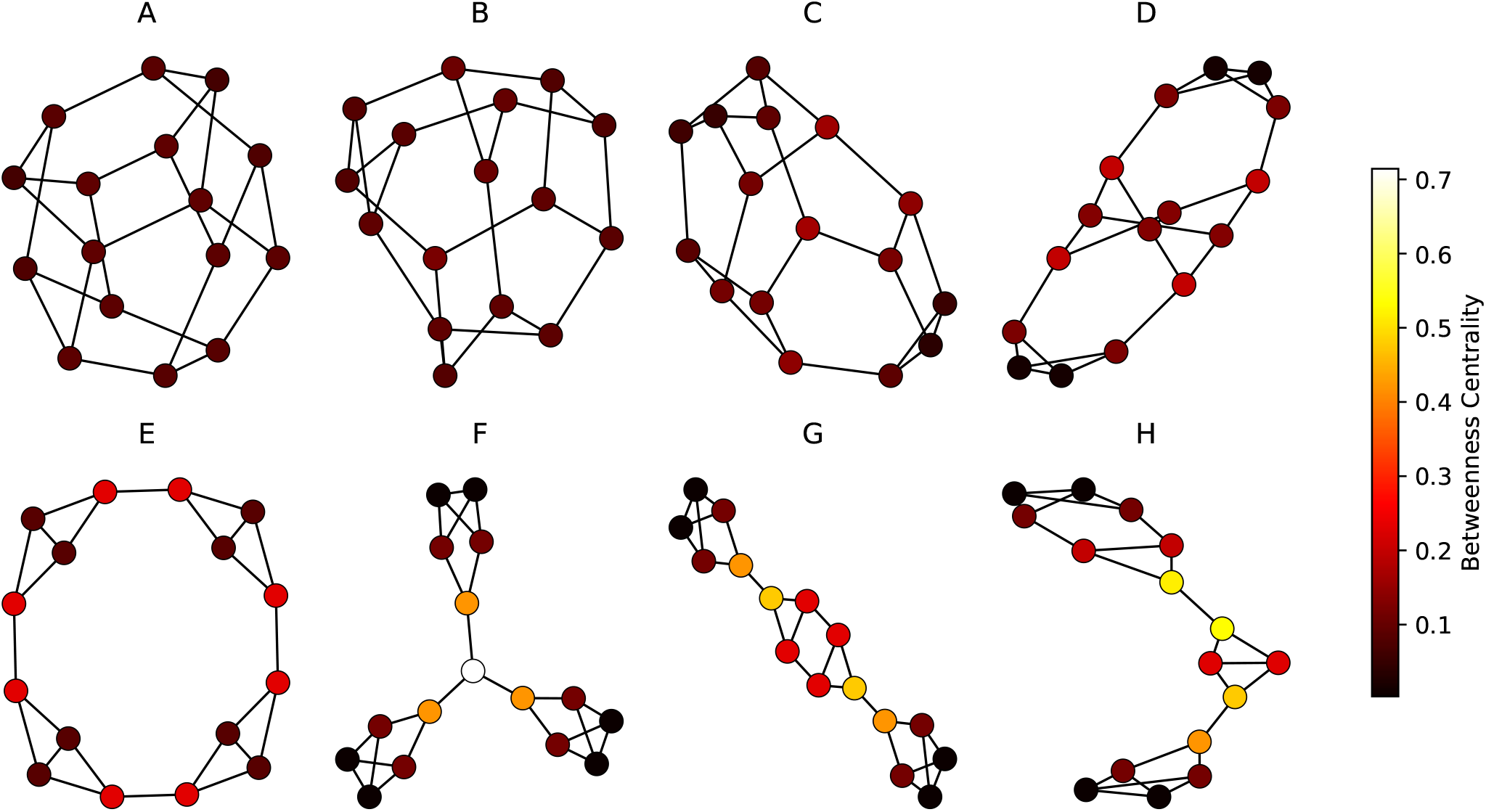
Eight network topologies with 16 nodes and 24 edges used to examine how communication patterns influence cumulative cultural evolution. Networks are ordered by average shortest path length and variation in betweenness centrality.

In spite of these structural variations, Figure 5 shows that no noticeable difference is observed among these eight networks in terms of the mean population complexity attained at equilibrium. In fact, we observe, in particular for direct bias, that structural properties play a role in how the equilibrium is approached, and thus, for intermediate timesteps, the less efficient networks *E* −*F* −*G*−*H* exhibit a transiently poorer performance.

**Figure 5.**
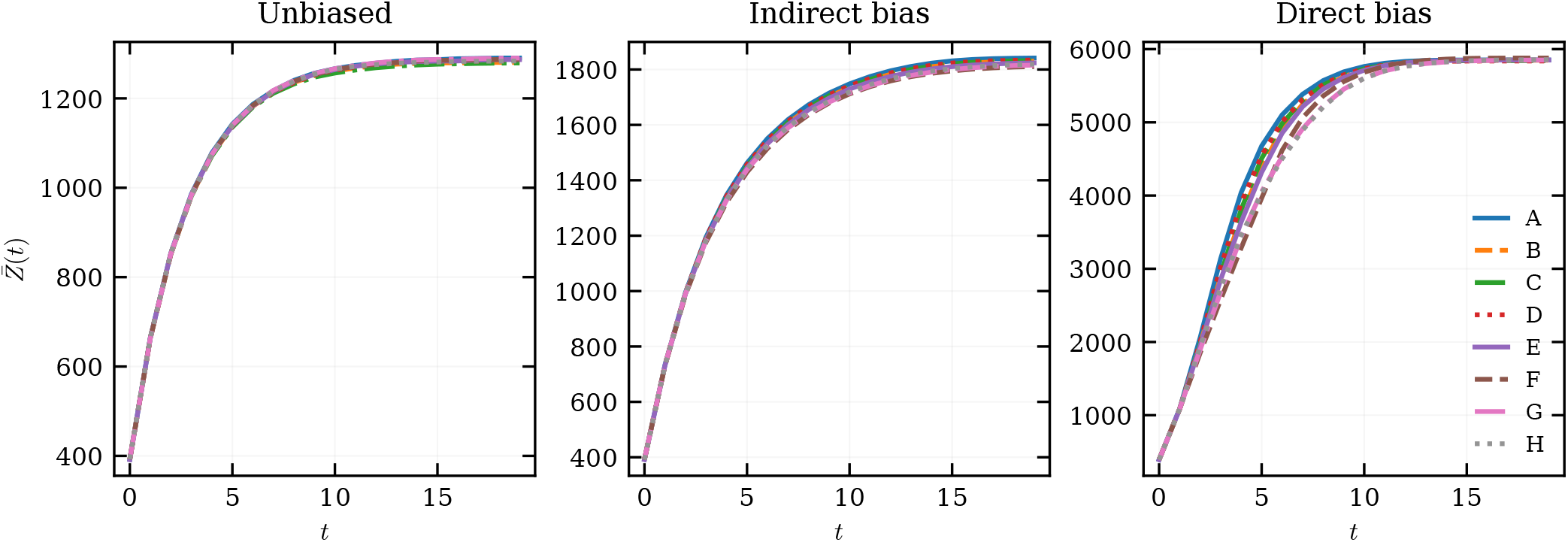
Temporal evolution of mean cultural complexity across the communication structures for each transmission rule. Parameter values are *λ* = 1000, *c*_*s*_ = 5, and *c*_*i*_ = 10.

### D. Time-dependent effort budget

To capture the idea that the constraints on cultural accumulation may relax over time, we consider a time-dependent effort budget rather than a fixed acquisition cost. This choice is closely related to the mechanism proposed by Mesoudi [17], who argues that CCE is limited because acquiring increasingly complex inherited knowledge becomes progressively more costly and time-consuming, leaving less time for innovation. In our formulation, however, the same trade-off can be represented from the opposite perspective: instead of letting acquisition costs rise or fall directly, we allow the effective effort available to individuals to vary in time. Mathematically, these two descriptions are equivalent up to a rescaling, since increasing the available budget has the same effect on net innovation capacity as reducing the cost of acquiring prior knowledge. In fact, technological innovations, institutional improvements, and accumulated knowledge can therefore expand the resources and capabilities available for learning and production [32].

We model improvements in learning and innovation capacity by allowing the individual effort budget to increase over timesteps. At the beginning of the process (*t* = 1), the budget is fixed at *λ*(0) = *λ*_0_, and in each subsequent timestep (*t*≥1), the effort budget increases monotonically toward an upper limit, *λ*_max_, according to

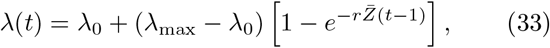

where *r* determines how rapidly the effort budget approaches its maximum value. In our simulations, we set λ_0_ = 100 and λ_max_ = 1000.

The analyses were conducted for three values of *r, r* = 0.1, 0.01, and 0.001. Figure 6 presents the time evolution of mean population complexity for the three transmission rules for an unstructured population. Interestingly, we note that, under the unbiased and direct bias rule, an increased rate *r* of budget effort increase leads to greater complexity at stationarity, whereas un-der the indirect bias rule, the opposite holds. Still, under the unbiased model, we observe that the values of complexity, 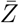, attained when *r* = 0.001 are considerably larger than those shown for an unstructured population in Figure 1, which uses the same set of parameters.

**Figure 6.**
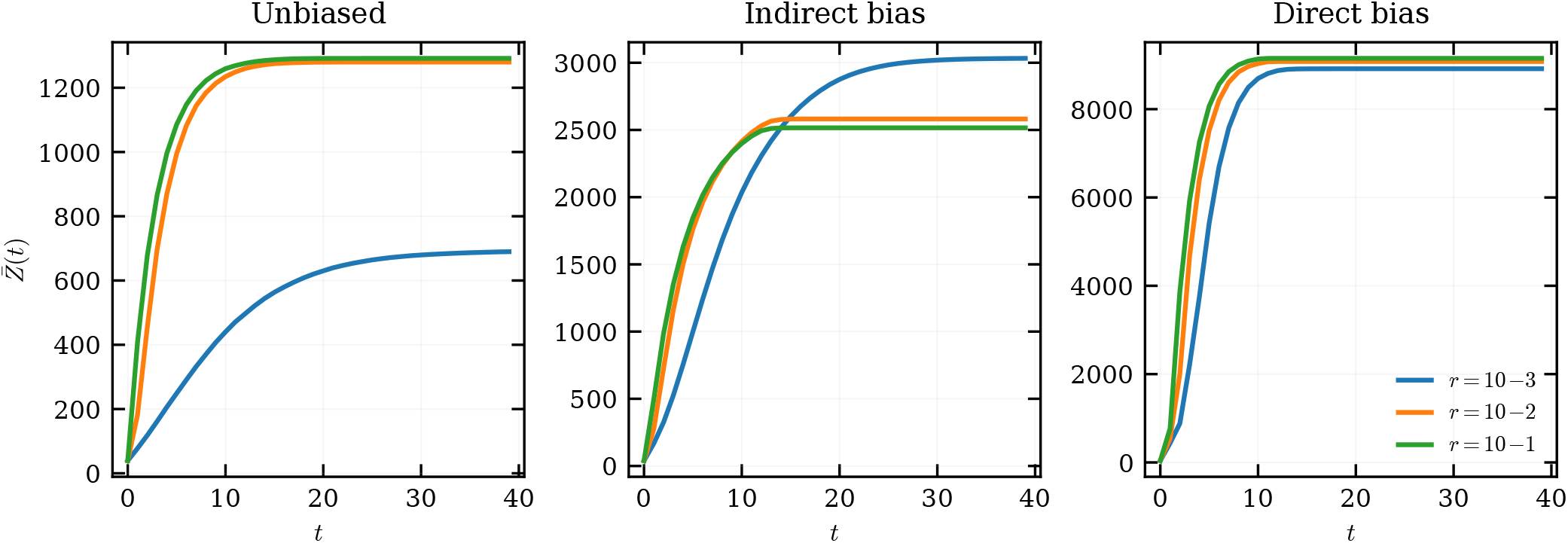
Temporal evolution of mean cultural complexity for different rates of budget increase, *r*, in homogeneous populations for each transmission rule. Parameter values are *N* = 100, *λ*_0_ = 100, *λ*_max_ = 1000, *c*_*s*_ = 5, and *c*_*i*_ = 10.

In Figure 7, we present simulation results for structured populations of fixed mean connectivity ⟨*k*⟩ = 10 under a time-dependent effort budget. We observe a qualitatively similar behavior to that verified for unstructured populations. A slight change is that, under direct bias, the mean population complexity collapses to a constant independent of *r* at stationarity for the three topologies. Figures A3 and A4 show results for *N* = 50 and *N* = 300, respectively. Qualitatively, we find, for *N* = 50, a first signature of the influence of the network topology: the mean population complexity at stationarity splits for scale-free networks under direct bias, with those evolving at a larger rate *r* performing better than those at low *r*.

**Figure 7.**
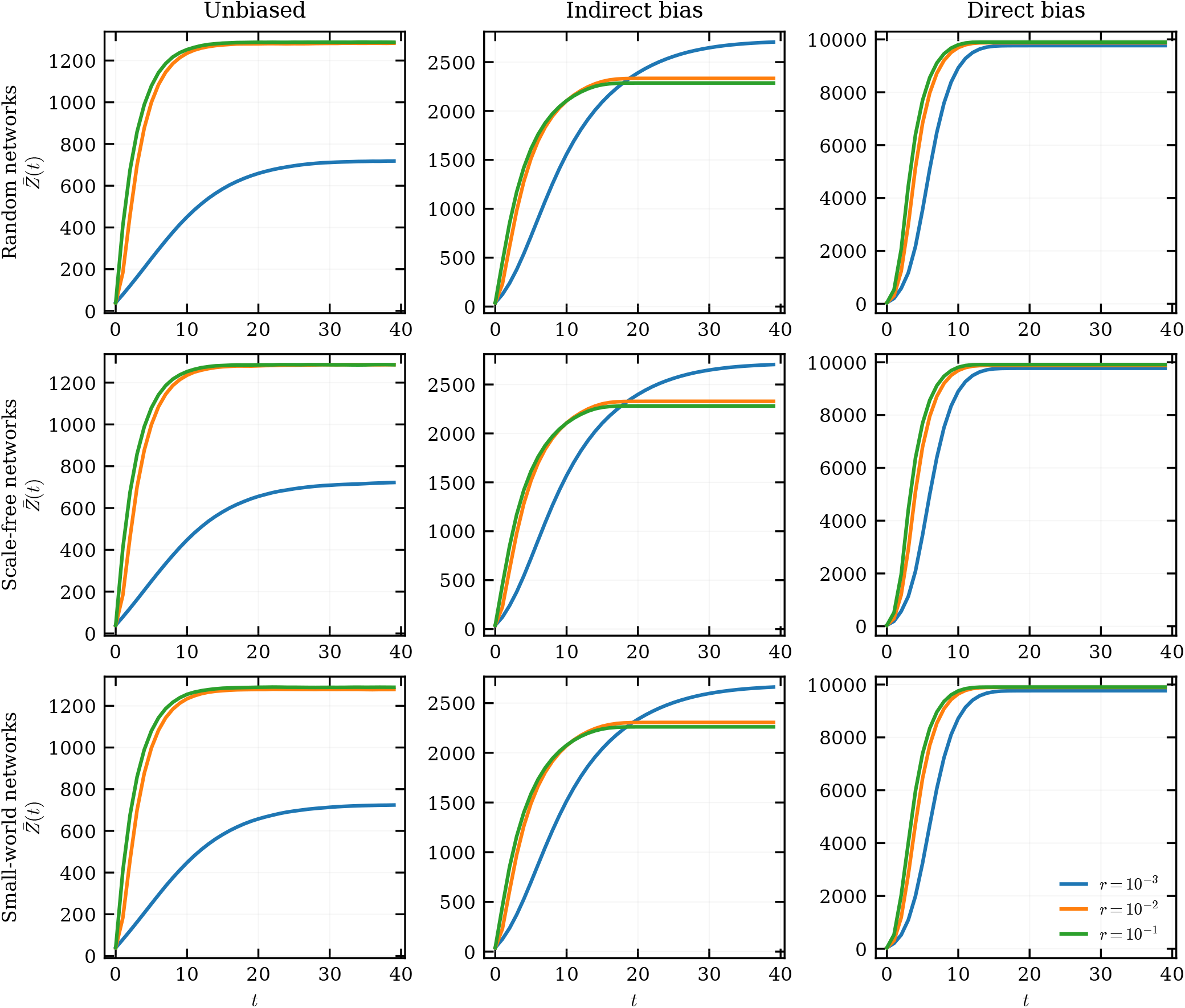
Temporal evolution of mean cultural complexity under a time-dependent effort budget for different rates of budget increase, *r*, in random, scale-free and small-world networks for each transmission rule. Parameter values are *N* = 100, ⟨*k*⟩ = 10, *λ*_0_ = 100, *λ*_max_ = 1000, *c*_*s*_ = 5, and *c*_*i*_ = 10.

For completeness, the same analysis is performed for the Mason-Watts networks (Figure A5). We observe no pronounced variation among the eight networks considered, indicating that the behavior remains consistent across all Mason-Watts instances studied.

## IV. DISCUSSION AND CONCLUSIONS

Our results show that the influence of social structure on CCE is strongly mediated by the mode of transmission. This is the main conclusion that emerges from the combination of the analytical treatment and the numerical simulations. In particular, the three learning rules considered here respond in markedly different ways to changes in connectedness and network organization. Under unbiased transmission, the dynamics are largely insensitive to both topology and mean degree, except in the low-connectivity regime, where unconnected components reduce the effective population size. Under indirect bias, increasing connectivity consistently enhances the cultural complexity attained at stationarity and accelerates convergence. Under direct bias, by contrast, the relationship is non-monotonic, with intermediate levels of connectivity maximizing cultural complexity. This pattern indicates that network effects cannot be discussed independently of the social-learning strategy through which information is sampled and transmitted.

The main implication of these findings is that connectedness emerges as the primary structural property of social architecture. Across all networks investigated in the present study, differences in stationary complex-ity (i.e. network topology) are comparatively modest relative to the effects of varying ⟨*k* ⟩ and, above all, the transmission rule itself. In other words, broad structural properties such as clustering, degree heterogeneity, and average shortest path length appear to play a secondary role relative to the simple fact of how many potential models are accessible and how learners use that information. This result is consistent with the idea that many topological effects may only become visible when the transmission process is itself sensitive to local variation, bottlenecks, or persistent lineage separation [33]. In our setting, such effects remain limited for most parameter combinations.

The analysis of the Mason–Watts communication networks reinforces this interpretation. These eight networks were designed to differ strongly in communication efficiency and centralization. Size and density were kept fixed. Yet, they produce little variation in final mean cultural complexity. Their main effect is instead transient, particularly under direct bias. In those cases, less efficient networks show slower intermediate dynamics but still approach similar stationary values. This suggests that, in the present model, some structural features are more important for the tempo of CCE than for its eventual ceiling. Put differently, communication architecture may shape how quickly populations approach equilibrium, even when it does not substantially alter equilibrium values.

Our findings reflect an important extension to previous work examining social networks and CCE. Cantor et al. [10] found similarly that no single network structure favored CCE under all conditions, as did we. However, we used a more elaborate model of CCE involving multiple hierarchical levels of cultural complexity, rather than just two semi-independent trait lineages, realistic time budgets which copying and innovation use up, and more elaborate transmission biases than the one-to-many or one-to-one transmission rules used in [10]. Indeed, our finding of systematic differences between transmission biases represents a valuable addition to this previous work, in which transmission (whether one-to-one or one-to-many) involved the unbiased random selection of demonstrators. We also found that the previously observed trade-off between speed of diffusion and maintenance of variation [16], leading to optimal CCE at intermediate levels of connectivity, only held for direct bias, and not unbiased or indirectly biased transmission. This again represents an important qualification to when we would expect to observe this trade-off in real cultural systems.

When transmission is unbiased and the population is uniform, our analytical derivations describe the longterm complexity with high accuracy. When we consider direct bias, the mathematical derivation improves further as the population size (*N*) increases, indicating that the approximation captures the large-*N* limit quite reliably. However, our mathematical formulation of the mean population complexity is less accurate for the indirect bias model. In that case, the analytical estimate tends to underestimate the actual complexity observed in the simulations. This makes sense, as the theoretical development does not fully account for how selection repeatedly *refines* repertoires that have already been optimized in earlier stages. Because of this gap, we can use the mathematical derivation as a lower-bound estimate.

Altogether, these findings contribute to the broader debate over whether there exist better social networks for driving cultural progress than those seen in contemporary societies. In line with recent research [10], our results suggest that there is no single gold-standard structure that works across the board. Instead, the impact of a social network depends heavily on how people are actually learning from one another. Therefore, we should not view social structure and learning rules as separate. They are deeply interconnected, and we must study their interactions to understand how culture evolves.

## Appendix A: Additional results

**Figure A1.**
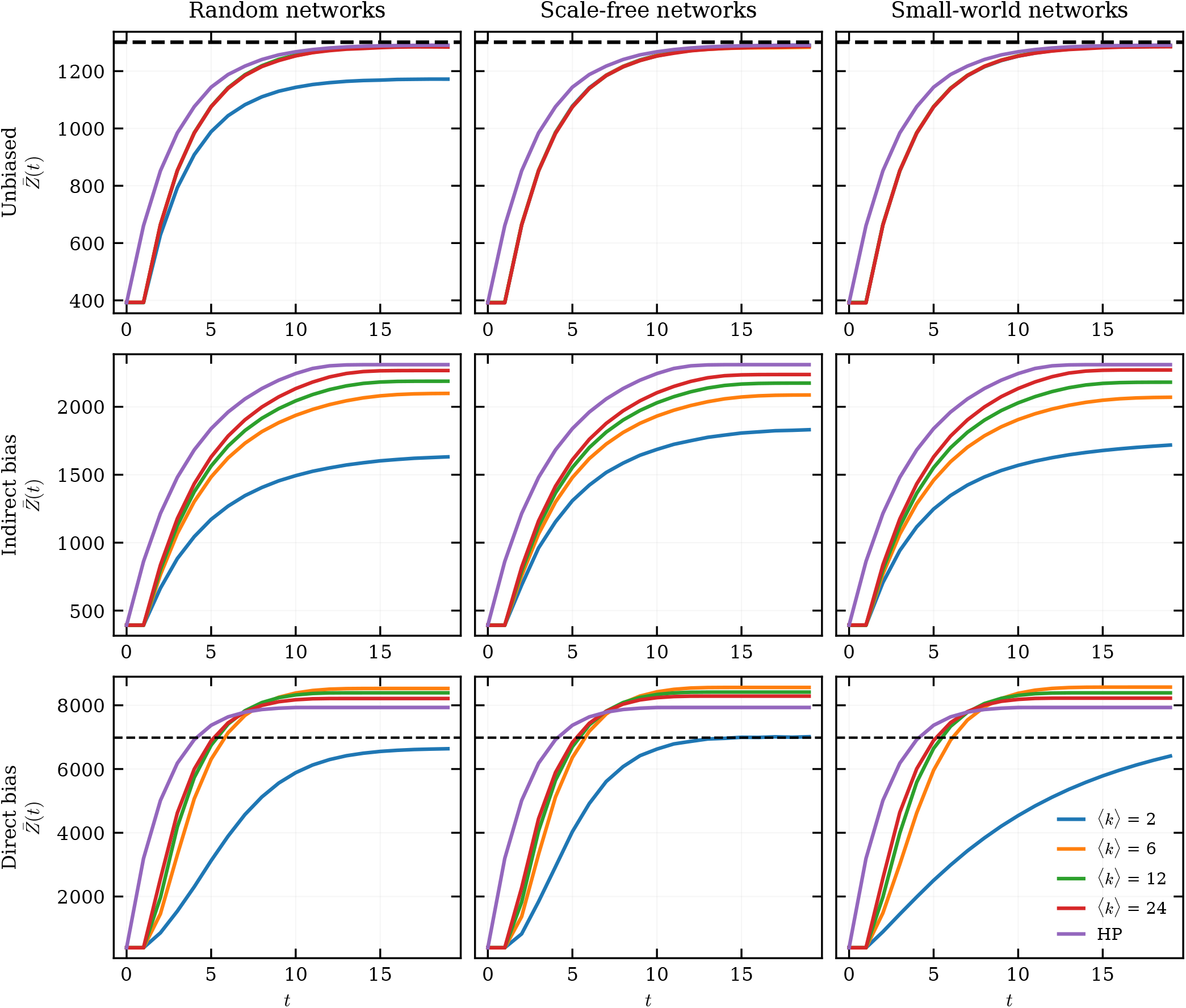
Temporal evolution of mean cultural complexity for representative values of average connectivity in random, scale-free, and small-world networks for each transmission rule. Parameter values are *N* = 50, *λ* = 1000, *c*_*s*_ = 5, and *c*_*i*_ = 10.

**Figure A2.**
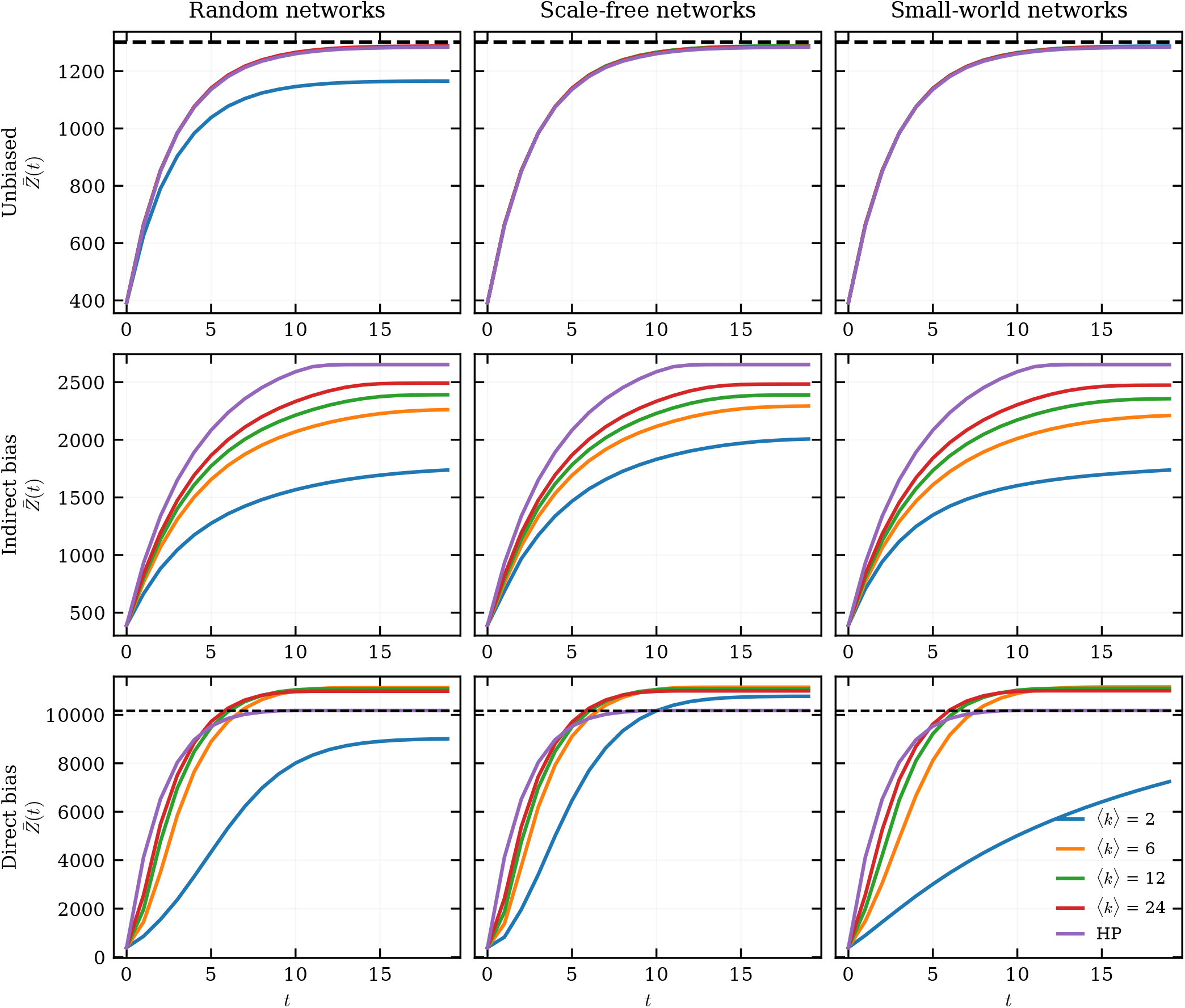
Temporal evolution of mean cultural complexity for representative values of average connectivity in random, scale-free, and small-world networks for each transmission rule. Parameter values are *N* = 300, *λ* = 1000, *c*_*s*_ = 5, and *c*_*i*_ = 10.

**Figure A3.**
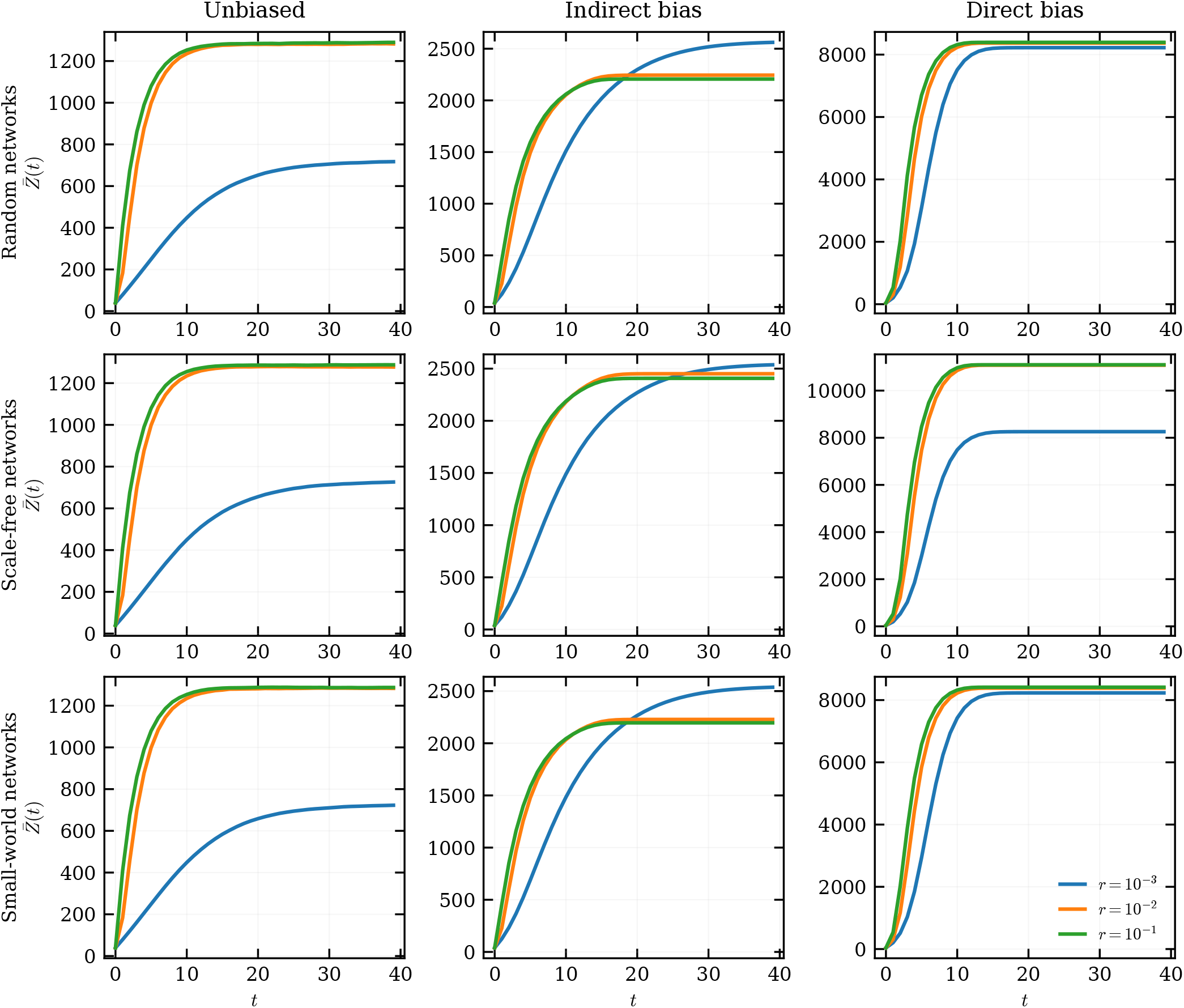
Temporal evolution of mean cultural complexity under a time-dependent effort budget for different *r* values in random, scale-free and small-world networks for each transmission rule. Parameter values are *N* = 50, ⟨*k* ⟩= 10, *λ*_0_ = 100, *λ* = 1000, *c*_*s*_ = 5, and *c*_*i*_ = 10.

**Figure A4.**
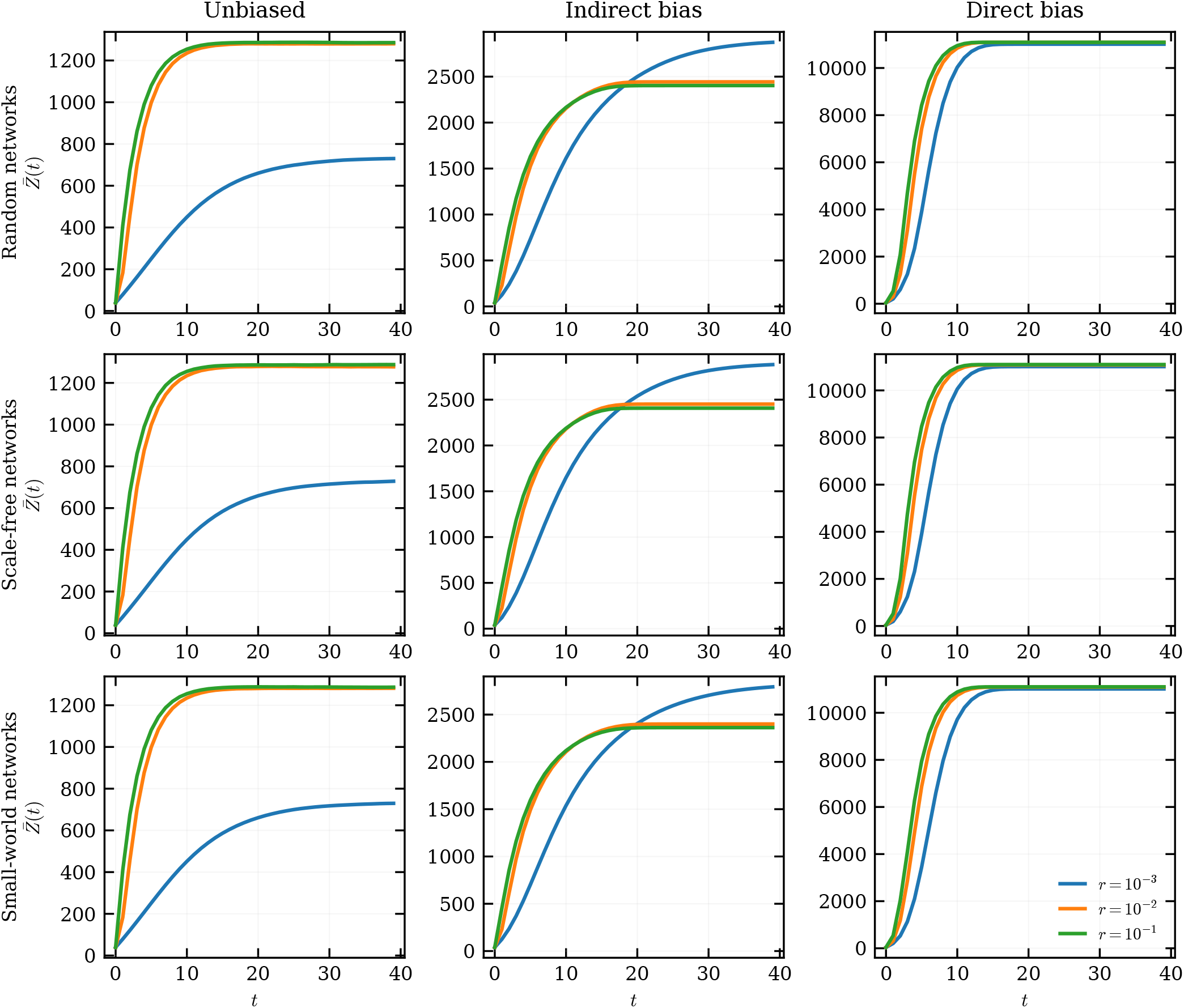
Temporal evolution of mean cultural complexity under a time-dependent effort budget for different *r* values in random, scale-free and small-world networks for each transmission rule. Parameter values are *N* = 300, ⟨*k*⟩= 10, *λ*_0_ = 100, *λ* = 1000, *c*_*s*_ = 5, and *c*_*i*_ = 10.

**Figure A5.**
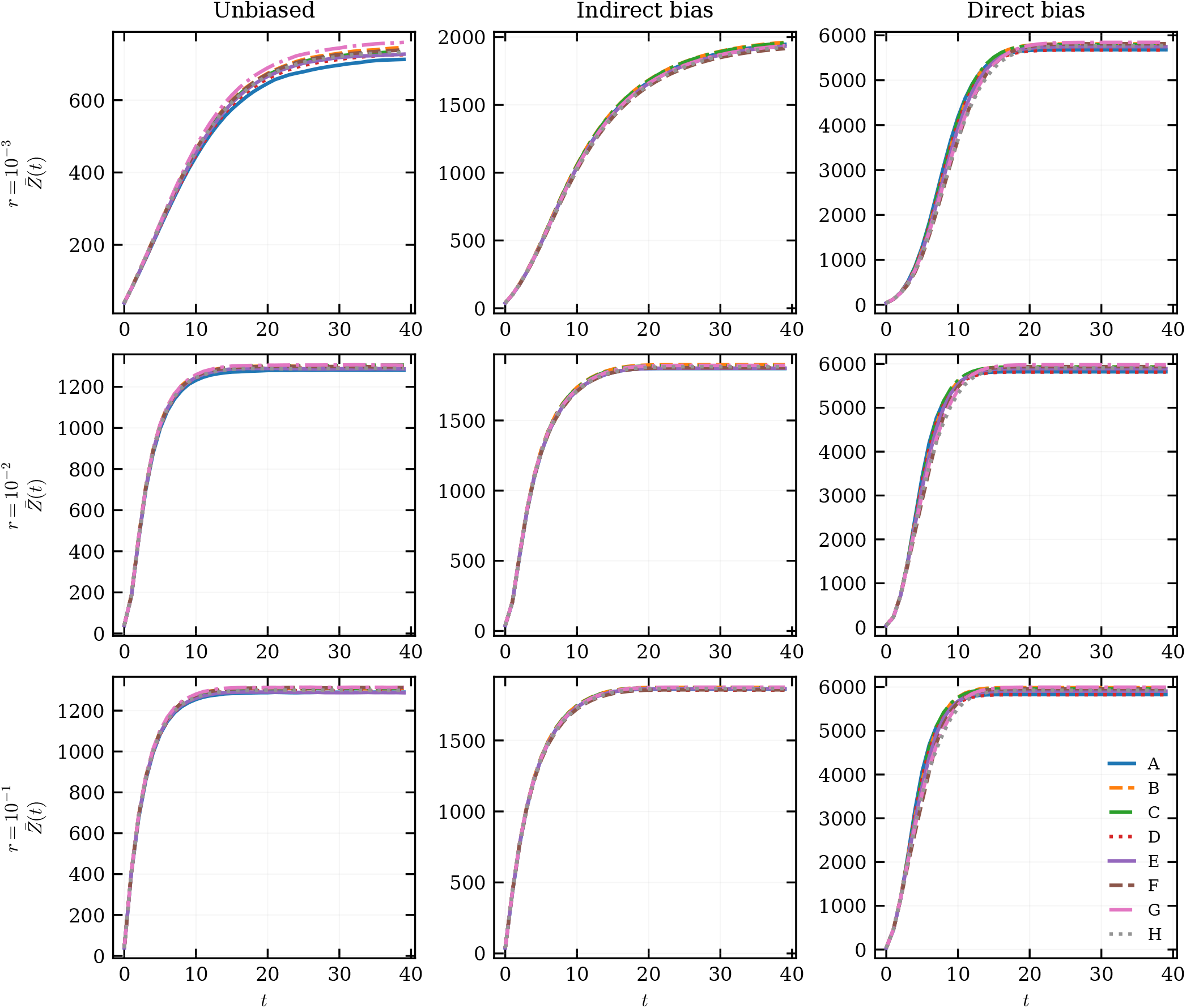
Temporal evolution of mean cultural complexity under a time-dependent effort budget for different *r* values across the communication structures for each transmission rule. Parameter values are *λ*_0_ = 100, *λ* = 1000, *c*_*s*_ = 5, and *c*_*i*_ = 10.

